# Shear stress magnitude affects bacterial extracellular vesicle stimulation of endothelial cells

**DOI:** 10.64898/2026.11.24.768965

**Authors:** Panteha Torabian, Abigail C. Wojehowski, Bradley Benedict, Owen B. Faust, Lea V. Michel, Thomas R. Gaborski

## Abstract

Extracellular vesicles released by pathogenic bacteria can induce strong inflammatory cascades, in part, by upregulating adhesion molecules on the surface of endothelial cells. The adhesion molecules support immune cell attachment and migration into surrounding tissues, amplifying local and systemic inflammation in conditions such as sepsis. We and others have predominantly studied the role of bacterial extracellular vesicles (BEVs) in tissue barrier models and microphysiological systems (MPS) under static conditions. Here, we investigated how physiological levels of fluid shear stress affect endothelial stimulation by Escherichia coli-derived BEVs. Human umbilical vein endothelial cells (HUVECs) were exposed to BEVs, lipopolysaccharide (LPS), or untreated medium under static, low-shear (1.5 dyn/cm^2^), and high-shear (10 dyn/cm^2^) conditions, while increases in the surface expression of intercellular adhesion molecule-1 (ICAM-1) were measured. BEVs elicited greater ICAM-1 expression than LPS under both low- and high-shear conditions, suggesting endothelial responses to BEVs depend on both the stimulus and mechanical environment. Although the highest absolute ICAM-1 expression occurred under high shear, the largest relative response to BEVs occurred under low shear. Together, these data show that hemodynamic shear stress is a critical determinant of BEV-associated endothelial inflammatory responses and a critical element for vascular MPS devices modeling conditions such as sepsis.

## INTRODUCTION

Sepsis is a life-threatening syndrome caused by a dysregulated host response to infection and remains a major cause of morbidity and mortality worldwide [1]. Much of its pathology comes from vascular endothelial dysfunction, which promotes leukocyte recruitment, loss of barrier integrity, dysregulated coagulation, uneven microvascular perfusion, and eventually organ injury [1]. The endothelium is an active interface where circulating microbial products, immune mediators, and hemodynamic forces converge [2]. One hallmark of endothelial activation is increased expression of intercellular adhesion molecule-1 (ICAM-1), which supports leukocyte adhesion and transendothelial migration and is an established marker of vascular inflammation [3].

Gram-negative bacteria release bacterial extracellular vesicles (BEVs), nanoscale lipid-bilayer particles that carry lipopolysaccharide (LPS), outer-membrane proteins, lipoproteins, nucleic acids, and other cargo [4]. Unlike intact bacteria, BEVs can spread bacterial signals beyond the site of infection and reach distant host cells, which may contribute to systemic inflammation. Our laboratory has shown that clinically relevant antibiotics change BEV production by *E. coli* [5], and that E. coli-derived BEVs induce dose-dependent endothelial ICAM-1 expression that is substantially reduced by inhibiting Toll-like receptor 4 (TLR4) [3]. A multicellular blood-brain barrier model later showed that interactions between endothelial and immune cells also shape BEV-associated vascular responses [6]. These studies established that BEVs can activate endothelium in static culture. How the response changes under flow remained unclear.

Blood flow is a major regulator of endothelial behavior. Wall shear stress varies along the vascular tree with vessel diameter, geometry, and local flow conditions [7], with lower values generally found in venous or low-flow environments and higher values in arteries [8]. It shapes endothelial morphology, cytoskeletal organization, barrier function, inflammatory signaling, and gene expression through shear-sensing structures that include the glycocalyx, junctional complexes, integrins, focal adhesions, and the cytoskeleton [9]. Endothelial cells therefore encounter circulating bacterial products in a flow-conditioned state, not the mechanically quiescent state of static culture. Vascular microphysiological systems (MPS) address this by perfusing endothelial cells at defined shear stresses [10]. Our group used a reconfigurable microfluidic µSiM platform to study endothelial barriers under flow and observed flow-induced alignment and changes in shear-responsive gene expression [11]. For studies of BEV-induced inflammation, this raises a question even before vesicles are introduced. How much does shear alone change endothelial ICAM-1 expression?

Flow may also influence how BEVs encounter the endothelium. BEVs and nanoparticles present LPS within a membrane that also contains other bacterial components [12]. Reaching the endothelium requires vesicle transport to the cell surface, followed by surface association, membrane interaction, or internalization [13], so the BEV concentration added to a system may differ from the effective dose at the cell surface, especially under fluid transport [14]. For mammalian extracellular vesicles and synthetic nanoparticles, endothelial association and uptake vary substantially with shear magnitude, particle properties, and endothelial state [9], [15], and shear stress can promote vesicle uptake [16]. Shear-dependent remodeling of the glycocalyx may also change how accessible the endothelial surface is to circulating particles [14], [19]. Identical bulk particle concentrations may therefore not produce equal endothelial association across flow conditions [17], [18], but this has not been established for bacterial vesicles.

Here, we used a microfluidic perfusion system to test how shear stress magnitude affects endothelial responses to *E. coli*-derived BEVs. HUVECs were exposed to BEVs, soluble LPS, or untreated medium under static, low-shear (1.5 dyn/cm^2^), and high-shear (10 dyn/cm^2^) conditions. These magnitudes were chosen to represent distinct, physiologically relevant mechanical environments, not a specific vascular bed [15], [17], [18]. Untreated cells defined the response to shear alone, and LPS served as a bacterial inflammatory control delivered in soluble form rather than on a vesicle [20]. We hypothesized that BEV-induced activation would vary with shear magnitude and measured ICAM-1 expression as the readout. High shear raised baseline ICAM-1, and BEVs produced greater ICAM-1 expression than LPS under both flow conditions. The highest absolute ICAM-1 expression occurred under high shear, but the largest relative response to BEVs occurred under low shear (about 6.8-fold over untreated cells, compared with 1.5-fold under high shear).

## MATERIALS AND METHODS

### Endothelial Cell Culture and Microfluidic Device Seeding

Pooled primary human umbilical vein endothelial cells (HUVECs) were obtained from ATCC (PCS-100-013), Lonza (C2519A), and Thermo Fisher Scientific (C01510C). Cells were maintained in Human Large Vessel Endothelial Cell Medium (Gibco, cat. no. M200500) supplemented with 2% Large Vessel Endothelial Supplement (Gibco, cat. no. A1460801) and 1% penicillin–streptomycin. Cultures were maintained at 37°C in a humidified atmosphere containing 5% CO₂, with medium replaced every 2–3 days. Cells were passaged at approximately 80% confluence and used for experiments at passages 7–8.

For microfluidic experiments, HUVECs were seeded into µ-Slide I 0.4 Luer microfluidic channels (ibidi GmbH, Martinsried, Germany), each consisting of a rectangular channel 50 mm in length, 5 mm in width, and 0.4 mm in height. Cells were seeded at a suspension concentration of approximately 5 × 10⁵ cells/mL, using 100 µL per channel. Following seeding, channels were maintained under static conditions for 24 h to allow cell attachment and formation of a confluent endothelial monolayer before the initiation of flow experiments.

### Isolation and Purification of *E. coli*-Derived BEVs

Escherichia coli (E. coli, strain CFT073) were cultured overnight on Lysogeny Broth (LB) agar at 37°C. Single colonies were inoculated into 50 mL of LB broth and incubated overnight at 37°C with shaking at 160 rpm. This primary culture was used to inoculate secondary LB cultures (200 mL each), which were grown to mid-logarithmic phase (OD_600_ ≈ 0.8). Cultures were subsequently incubated at 37°C with shaking at 180 rpm for an additional 3 h. To isolate BEVs, bacterial cells were pelleted by centrifugation at 5,000 × *g* for 15 min at 4°C. The resulting supernatant was collected and passed through a 0.22 µm polyethersulfone (PES) syringe filter to remove residual bacterial cells and larger debris. The clarified filtrate was subsequently ultracentrifuged at 100,000 × *g* for 1 h at 4°C to pellet the BEVs. After removal of the supernatant, the BEV pellets were resuspended in sterile, endotoxin-free phosphate-buffered saline (PBS).

To assess residual viable bacterial contamination, aliquots of the isolated BEV preparations were plated onto LB agar and incubated overnight at 37°C. No visible bacterial colonies were observed. BEV preparations were stored at 4°C for immediate use or at −20°C until subsequent experimental analysis.

### Western Blot Analysis

Western blotting was performed to detect the outer membrane proteins OmpA and peptidoglycan-associated lipoprotein (Pal) in isolated *E. coli*-derived BEVs. Samples were separated using standard SDS-PAGE (4–16% bis-acrylamide precast gels, Avantor), with equal sample volumes loaded into each well. For immunoblotting, rabbit polyclonal anti-outer membrane protein A (OmpA; approximately 38 kDa; Antibody Research Corporation, St. Peters, MO, USA) and mouse anti-Pal antiserum raised against purified recombinant non-lipidated Pal protein (approximately 21 kDa, containing a 6×His tag for purification; Rochester General Hospital) were used, as previously described [5]. To confirm the presence of outer membrane components in BEV preparations, we probed for OmpA and Pal, two abundant outer membrane-associated proteins.

Immunoblots were developed using SuperSignal West Femto Maximum Sensitivity Substrate (Thermo Fisher Scientific, Waltham, MA, USA) and imaged using a Bio-Rad ChemiDoc Imaging System.

### Transmission Electron Microscopy (TEM)

Negative-stain transmission electron microscopy (TEM) was performed to examine the morphology of isolated E. coli-derived BEVs. Formvar/carbon-coated 200-mesh copper grids were glow-discharged using a PELCO easiGlow system for 30 s at 30 mA. Subsequently, 3 µL of BEV suspension was applied to each grid and allowed to adsorb for 30 s. Excess liquid was removed using filter paper, and the grids were washed three times with 15 µL of molecular-grade water.

Negative staining was performed by applying 10 µL of filtered 0.75% uranyl formate solution twice, with excess liquid removed using Whatman Grade 50 filter paper between applications. Following air drying, the grids were examined using a Talos 120C transmission electron microscope (Thermo Fisher Scientific). Images were acquired using a CETA 16-megapixel camera and TIA imaging software (Thermo Fisher Scientific).

### Microfluidic Perfusion System and Hydrodynamic Characterization

A closed-loop microfluidic perfusion system was assembled to expose confluent HUVEC monolayers to controlled fluid shear stress. The system consisted of an ibidi µ-Slide I 0.4 Luer microfluidic channel connected to a peristaltic pump (Masterflex L/S Easy-Load, Cole-Parmer, Vernon Hills, IL, USA), medium reservoirs, and silicone tubing. The medium reservoirs were incorporated into the system to dampen the flow pulsations generated by the peristaltic pump, maintaining a more consistent flow rate and shear stress within the microfluidic channel. The assembled perfusion circuit was maintained within a humidified cell culture incubator at 37°C under 5% CO₂ throughout the experimental period.

The rectangular microfluidic channels had an internal width of 5 mm and height of 0.4 mm. Wall shear stress was estimated from the volumetric flow rate, medium viscosity, and channel dimensions using the approximation for fully developed laminar flow between parallel plates: τ= η × 131.6 × Φ. Where τ is wall shear stress (dyn/cm^2^), η is dynamic viscosity of the medium, and Φ is volumetric flow rate (mL/min). Based on this relationship, volumetric flow rates were continuously driven at either 1.56 mL/min or 10.55 mL/min to generate nominal wall shear stresses of 1.5 dyn/cm^2^ (low flow rate, LFR) and 10 dyn/cm^2^ (high flow rate, HFR), respectively.

To assess whether the experimental conditions were consistent with laminar, fully developed flow within the imaging region, hydrodynamic parameters were calculated for the rectangular channel geometry (length L = 50 mm, width w = 5 mm, height h = 0.4 mm). The hydraulic diameter (Dh) of the cross-section was calculated as: Dh = 4A / P = 2wh / (w + h) = 2(5 mm)(0.4 mm) / (5 mm + 0.4 mm) = 0.741 mm (7.41 × 10^-4 m). At 37°C, the cell culture medium was modeled with dynamic viscosity eta = 0.74 × 10^-3 Pa·s and density rho = 1000 kg/m³. Mean fluid velocities corresponded to 0.013 m/s at the low flow rate (1.56 mL/min; 1.5 dyn/cm^2^) and 0.088 m/s at the high flow rate (10.55 mL/min; 10 dyn/cm^2^). Corresponding Reynolds numbers (Re = rho * u_m * Dh / eta) were 13.0 for low shear and 88.1 for high shear, supporting the assumption of laminar flow (Re << 2300) across all experimental runs. The hydrodynamic entrance length (Le) required for the velocity boundary layers to coalesce and approach a fully developed velocity profile was calculated using the laminar duct correlation: Le = 0.05 * Re * Dh. Under the maximum operational flow rate (10.55 mL/min, Re = 88.1), the entrance length was 3.26 mm. Because quantitative epifluorescence imaging was performed along the channel centerline within the central 20 mm region, fields of view were positioned beyond the estimated hydrodynamic entrance length and away from the lateral boundaries, minimizing potential inlet and sidewall effects. These conditions support the use of nominal wall shear stress estimates within the imaging region.

### Shear Preconditioning and Host-Pathogen Experimental Design

After 24 h of static attachment, flow-conditioned endothelial monolayers were preconditioned at 1.5 dyn/cm^2^ for 24 h before treatment. Cells were then exposed for an additional 24 h to vehicle control, *E. coli*-derived BEVs (1 × 10⁸ particles/mL), or LPS (10 ng/mL; Sigma-Aldrich, cat. no. L3012-5MG) under their assigned static, low-shear (1.5 dyn/cm^2^), or high-shear (10 dyn/cm^2^) conditions. Treatments were introduced into the recirculating medium for the flow groups. Three independent microfluidic chips were used per condition (n = 3).

### Nanoparticle Tracking Analysis (NTA)

Nanoparticle size distributions and particle concentrations were assessed using a NanoSight NS300 system (Malvern Panalytical, Malvern, UK) equipped with an sCMOS camera, 532 nm laser, and 565 nm long-pass filter. BEV preparations were diluted in PBS to obtain an appropriate tracking density (approximately 1 × 10^8−1^ × 10^9^ particles/mL). Three 30-s videos were acquired per sample using camera level 15 and detection threshold 5. Hydrodynamic diameter was estimated from Brownian motion, and particle concentrations were corrected for sample dilution.

### Immunofluorescence Staining and ICAM-1 Morphometry

Following the 24 h treatment period, microfluidic channels were disconnected from the perfusion system and incubated with mouse anti-human ICAM-1 (CD54, clone HA58) primary antibody diluted 1:100 in M200 medium for 15 min at 37°C. Following primary antibody incubation, channels were washed three times with phosphate-buffered saline (PBS) and fixed in situ with 3.7% formaldehyde for 15 min at room temperature. Endothelial monolayers were subsequently washed with PBS and blocked with 10% normal goat serum (Thermo Fisher Scientific, cat. no. 50062Z) for 30 min at room temperature to minimize nonspecific antibody binding. Cells were then incubated with Alexa Fluor 488-conjugated goat anti-mouse IgG secondary antibody (Thermo Fisher Scientific, cat. no. A-11001), diluted 1:200 in blocking buffer, for 2 h at room temperature protected from light. Cell nuclei were counterstained with DAPI (Thermo Fisher Scientific, cat. no. D1306), diluted 1:400 in deionized water, for 3 min at room temperature, followed by three PBS washes.

Epifluorescence images were acquired using consistent exposure times, illumination intensities, and detector settings across experimental conditions to enable quantitative comparisons of ICAM-1 fluorescence. Three fields of view were acquired along the central region of each microfluidic channel, avoiding the lateral boundaries to minimize potential edge-related effects.

### Digital Image Processing and Quantitative Fluorescence Analysis

Epifluorescence micrographs were processed using a custom Python-based image analysis workflow for fluorescence thresholding, channel merging, and image format conversion. To ensure consistency across experimental conditions, fixed intensity thresholds (111 and 1108) were applied to the green fluorescent protein (GFP)/ICAM-1 fluorescence channel, as appropriate for the corresponding image acquisition settings, whereas DAPI channel thresholds were adjusted to account for variations in nuclear fluorescence intensity.

Quantitative image analysis was performed using QuPath (version 0.7.0), an open-source software platform for digital image analysis [21]. A custom Groovy script was implemented to automate cell detection, fluorescence intensity measurements, and extraction of 13 Haralick texture features. Cell detection was performed using DAPI nuclear staining and QuPath’s standard detection parameters, with watershed-based segmentation used to distinguish adjacent nuclei. Detected cell populations were used to quantify cell numbers and fluorescence intensity measurements across experimental conditions.

To evaluate the consistency of automated cell detection, representative images were independently analyzed using Fiji/ImageJ, and nuclear counts were compared with those obtained using QuPath. Preliminary parameter optimization included evaluation of nuclear detection thresholds, minimum object area, and Gaussian smoothing. Following this evaluation, standardized detection settings were applied consistently across experimental images.

For experiments investigating CellMask Orange (CMO)-labeled bacterial extracellular vesicles (BEVs), fluorescence micrographs were processed using a uniform fluorescence intensity threshold across the complete dataset. DAPI-positive nuclei were used to determine cell numbers, while CMO-associated fluorescence (red fluorescent protein /RFP,channel) and ICAM-1 expression (GFP channel) were quantified for each image. Representative measurements were independently cross-validated using Fiji/ImageJ to assess consistency between image analysis methods.

### Statistical Analysis

Statistical analyses were performed using GraphPad Prism (version 10; GraphPad Software, Boston, MA, USA). Three independent microfluidic chips were evaluated per experimental condition (n = 3), and three imaging fields (inlet, middle, and outlet regions) were measured within each chip. For primary ICAM-1 comparisons, the three field-level measurements were averaged to obtain one mean value per chip. The three chip means per condition were plotted individually with group mean ± standard deviation (SD) in **Fig. 3A–C**. Differences among static, LFR, and HFR conditions within each treatment group, as well as comparisons across all nine treatment–shear conditions, were evaluated by ordinary one-way ANOVA followed by Tukey’s multiple-comparisons test using chip means as independent observations. In **Fig. 3E**, all nine imaging-field measurements per condition were displayed to illustrate within-chip and spatial variation; these fields were not treated as independent biological replicates for the primary ICAM-1 significance tests.

Data distributions were assessed using the Shapiro–Wilk and D’Agostino–Pearson normality tests as appropriate. For CellMask Orange (CMO) fluorescence analysis, three imaging-field measurements were averaged to obtain one mean value per independent chip (n = 3 chips per shear condition). Differences in CMO fluorescence among static, LFR, and HFR conditions were evaluated using an exact Kruskal–Wallis test followed by Dunn’s multiple-comparisons test. The relationship between CMO and ICAM-1 fluorescence was evaluated using paired chip-level means from the same experimental chips (n = 9 across three shear conditions) by ordinary least-squares linear regression. Statistical significance was defined as p < 0.05, with adjusted p-values used for multiple comparisons.

### Fluorescent Labeling of Bacterial Extracellular Vesicles

Bacterial extracellular vesicles (BEVs) were fluorescently labeled using CellMask Orange (CMO; Thermo Fisher Scientific), a lipophilic fluorescent membrane dye. Prior to labeling, BEV concentrations were determined using nanoparticle tracking analysis (NTA), and samples were diluted to 4 × 10⁸ particles/mL in 0.22 µm-filtered phosphate-buffered saline (PBS).

A CMO working solution was prepared through sequential dilutions of the original stock in PBS (1:20, followed by 1:10 and 1:5), resulting in a 1:1,000 dilution relative to the stock dye. Subsequently, 5 µL of the diluted CMO working solution was added to 500 µL of BEV suspension, yielding an approximate final dye dilution of 1:101,000 relative to the original stock. Samples were protected from light and incubated at room temperature for 45 min to facilitate membrane labeling. Following fluorescent labeling, BEV preparations were diluted to a final concentration of 1 × 10^8^ particles/mL before endothelial exposure.

Fluorescent labeling was evaluated using fluorescence-mode nanoparticle tracking analysis (F-NTA) with a 562 nm optical filter. Corresponding scatter-mode NTA measurements were performed to determine total particle concentrations. The proportion of fluorescently detectable particles was estimated by comparing fluorescence-mode and scatter-mode particle concentrations. CMO-labeled BEVs were subsequently used in endothelial association experiments, with CMO-associated fluorescence quantified using epifluorescence microscopy.

## RESULTS

### Characterization of isolated *E. coli*-derived BEVs

*E. coli*-derived bacterial extracellular vesicles (BEVs) were characterized using nanoparticle tracking analysis (NTA), transmission electron microscopy (TEM), and Western blotting to evaluate particle size, morphology, and the presence of bacterial outer membrane-associated proteins **(Fig. 1)**.

**Figure 1.**
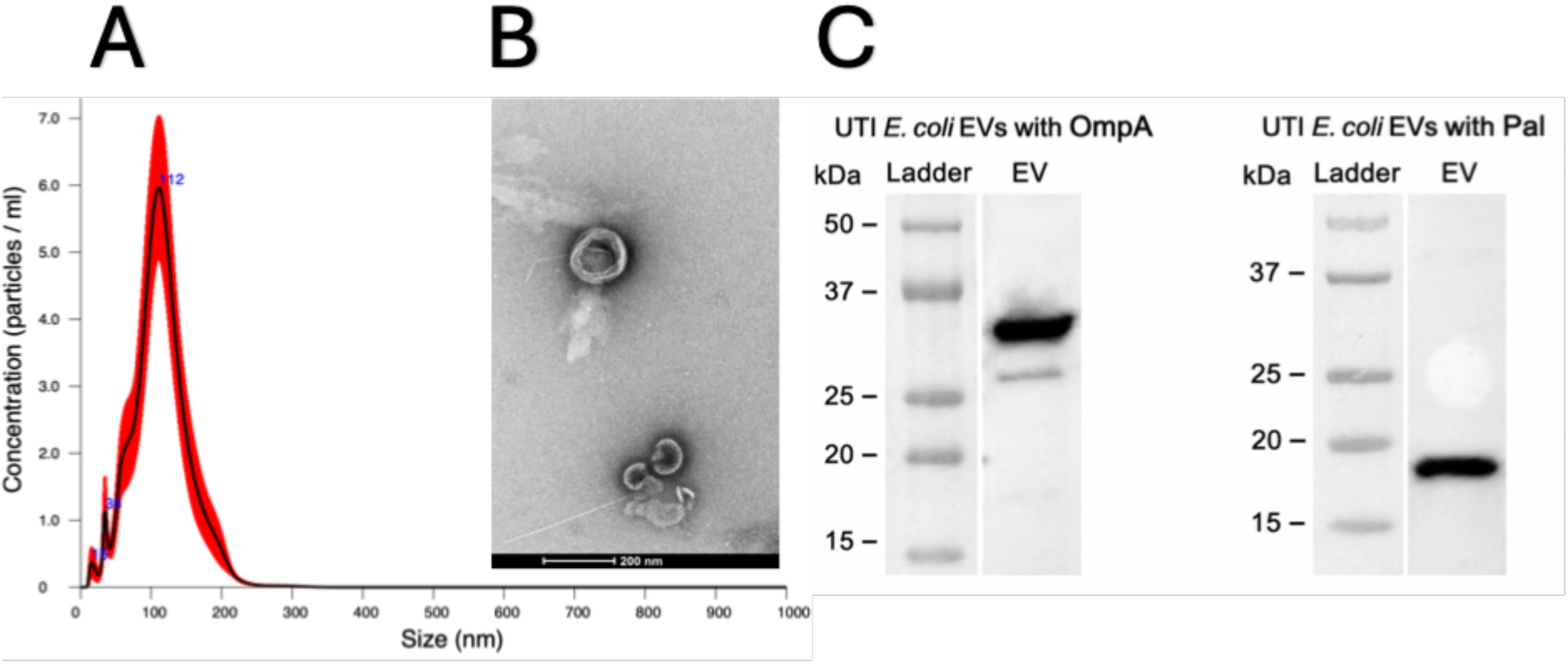
Physicochemical and molecular characterization of E. coli -derived bacterial extracellular vesicles. **(A)** Representative nanoparticle tracking analysis (NTA) showing the particle size distribution of isolated BEVs, with a modal diameter of approximately 112 nm. **(B)** Representative negative-stain transmission electron microscopy (TEM) image showing vesicle-like structures (scale bar = 200 nm). **(C)** Representative Western blots showing detection of the outer membrane protein A (OmpA; ∼35 kDa) and peptidoglycan-associated lipoprotein (Pal; ∼18 kDa) in isolated BEV preparations. Molecular weight markers (kDa) are shown alongside the samples.

NTA revealed a predominant particle population with a modal hydrodynamic diameter of approximately 112 nm **(Fig. 1A)**. Negative-stain TEM imaging demonstrated nanoscale vesicle-like structures with rounded and cup-shaped appearances, consistent with the expected morphology of isolated BEVs **(Fig. 1B)**. Western blot analysis detected outer membrane protein A (OmpA; approximately 35 kDa) and peptidoglycan-associated lipoprotein (Pal; approximately 18 kDa), supporting the presence of bacterial outer membrane-associated proteins in the isolated BEV preparations **(Fig. 1C)**.

Together, these results support the successful isolation and characterization of *E. coli*-derived BEVs used in subsequent endothelial stimulation experiments.

### Microfluidic perfusion established distinct low- and high-shear endothelial environments

A closed-loop microfluidic perfusion system was established to investigate how defined fluid shear stresses influence endothelial responses to *E. coli*-derived BEVs **(Fig. 2A)**. HUVECs were cultured within ibidi µ-Slide I 0.4 Luer microfluidic channels and exposed to volumetric flow rates of 1.56 and 10.55 mL/min, corresponding to nominal wall shear stresses of 1.5 dyn/cm^2^ (low flow rate, LFR) and 10 dyn/cm^2^ (high flow rate, HFR), respectively. Wall shear stress was estimated using the rectangular-channel relationship illustrated in **Fig. 2B**, which also shows the corresponding equation for an idealized circular blood vessel for comparison.

**Figure 2.**
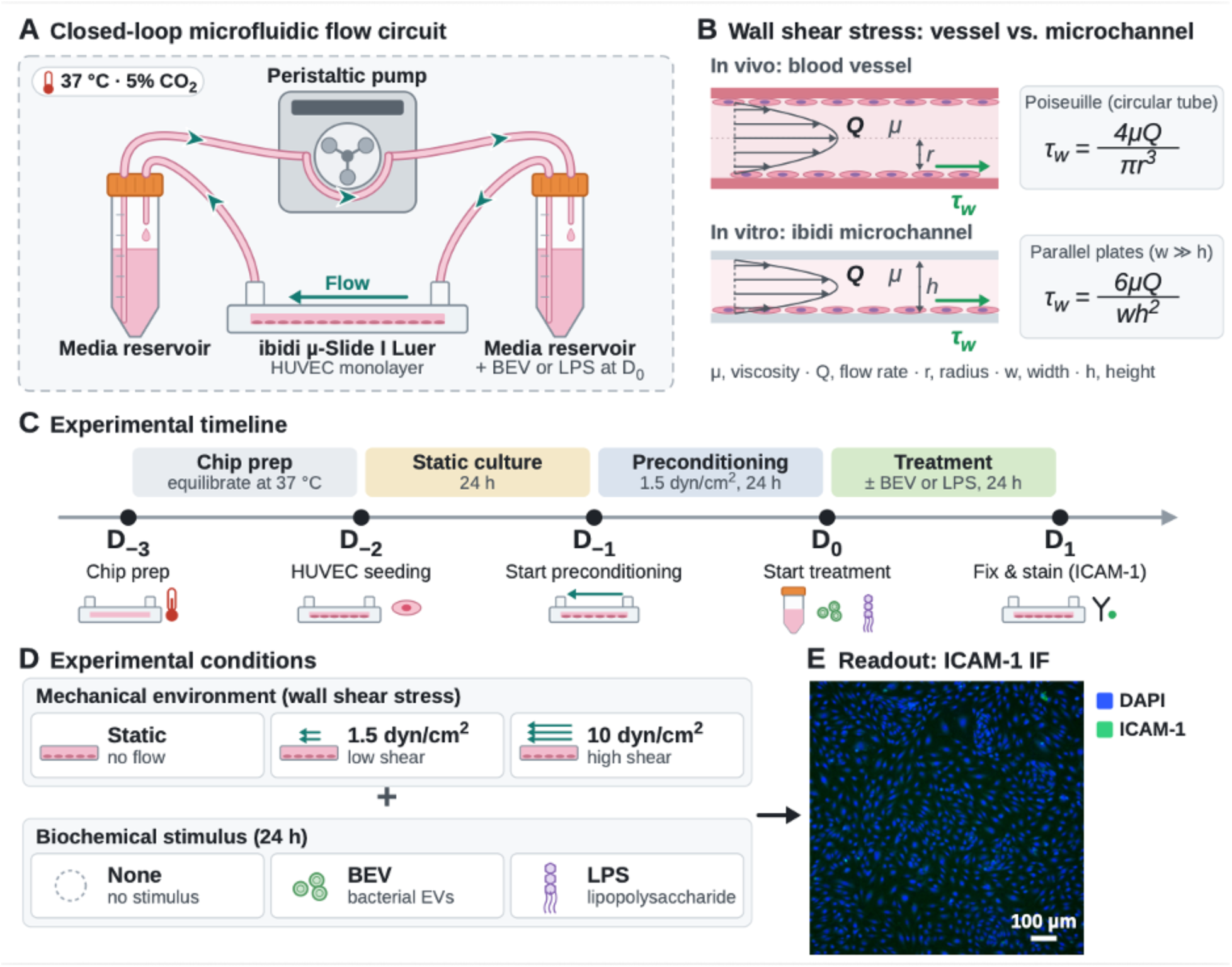
Experimental design and microfluidic perfusion system for investigating endothelial responses to bacterial extracellular vesicles under defined shear stress. **(A)** Schematic of the closed-loop microfluidic perfusion system consisting of a peristaltic pump, medium reservoirs, and an ibidi µ-Slide I 0.4 Luer channel containing a HUVEC monolayer. The system was maintained at 37 °C and 5% CO₂ throughout perfusion. **(B)** Comparison of idealized wall shear stress relationships for laminar flow in a circular blood vessel and a rectangular microchannel. The circular-tube equation is shown for physiological context, whereas experimental shear stress was estimated using the geometry-specific relationship for the rectangular ibidi channel. **(C)** Experimental timeline showing microfluidic channel preparation, HUVEC seeding and static attachment for 24 h, followed by 24 h of preconditioning at 1.5 dyn/cm^2^ for flow-conditioned groups only. Endothelial monolayers were subsequently exposed to their assigned treatments for an additional 24 h. **(D)** Experimental conditions combining three mechanical environments—static culture (0 dyn/cm^2^), low shear (1.5 dyn/cm^2^), and high shear (10 dyn/cm^2^)—with three biochemical treatments: untreated medium, E. coli-derived bacterial extracellular vesicles (BEVs; 1 × 10^8^ particles/mL), or lipopolysaccharide (LPS; 10 ng/mL). **(E)** Representative immunofluorescence micrograph of a HUVEC monolayer showing intercellular adhesion molecule-1 (ICAM-1; green) and DAPI-stained nuclei (blue). Scale bar = 100 µm.

Gravimetric flow measurements were consistent with the calculated flow-rate targets for both conditions. Differential pressure monitoring also demonstrated an increase in pressure drop with increasing flow rate, consistent with the expected behavior of the perfusion system. Together, these measurements supported the ability of the system to establish distinct and controlled flow conditions.

Following initial cell attachment, flow-conditioned HUVEC monolayers were preconditioned at 1.5 dyn/cm^2^ for 24 h before experimental treatment **(Fig. 2C)**. Cells were subsequently exposed to untreated control medium, BEVs (1 × 10⁸ particles/mL), or lipopolysaccharide (LPS; 10 ng/mL) for an additional 24 h under their assigned static, LFR, or HFR conditions **(Fig. 2D)**. Endothelial activation was evaluated by ICAM-1 immunofluorescence **(Fig. 2E)**.

### High shear stress increased basal endothelial ICAM-1 expression

To determine how shear stress independently influences endothelial activation, ICAM-1 fluorescence was quantified in untreated HUVECs maintained under static, LFR (1.5 dyn/cm^2^), and HFR (10 dyn/cm^2^) conditions **(Fig. 3A,D)**.

**Figure 3.**
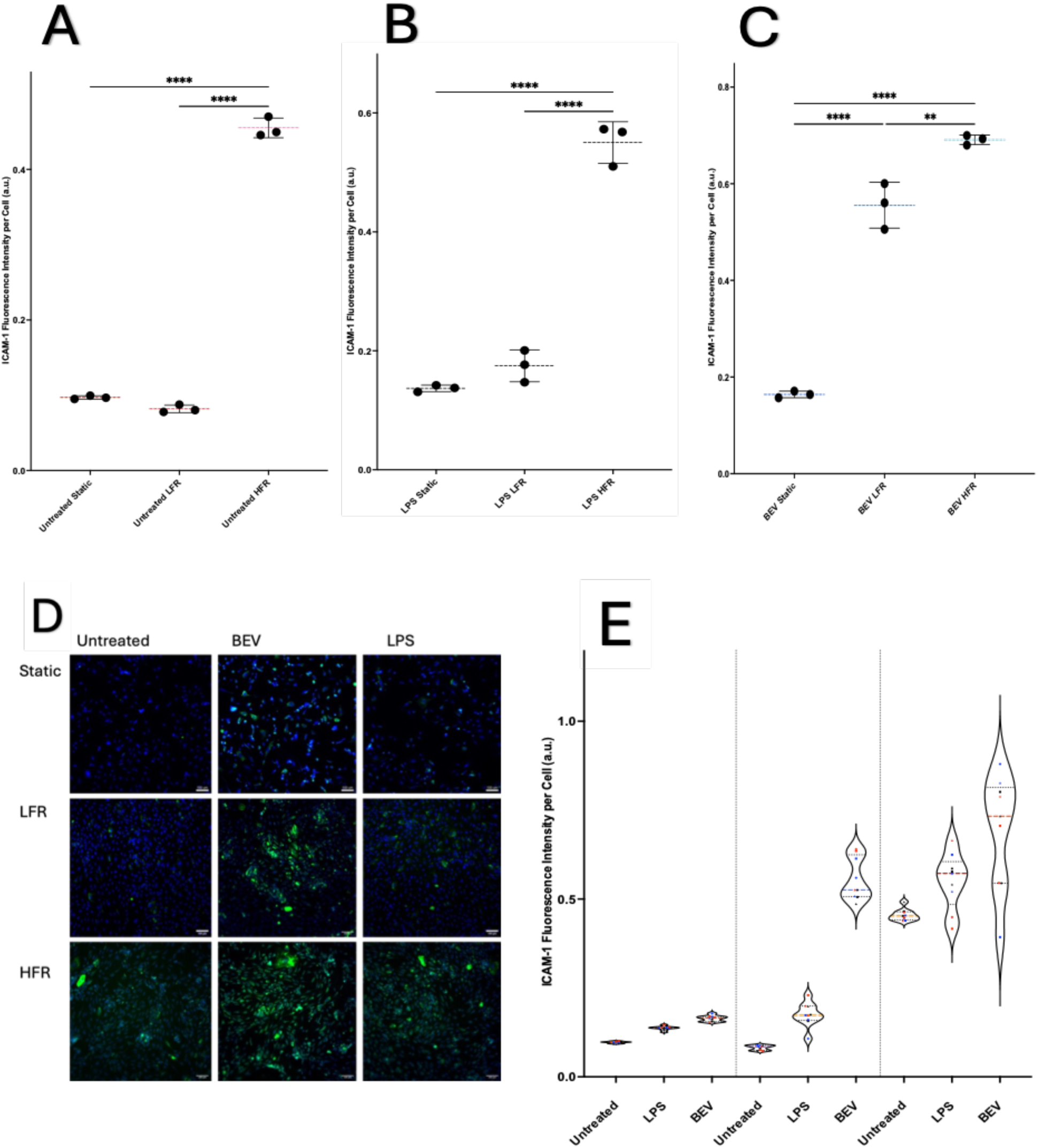
Shear-dependent ICAM-1 expression in untreated, LPS-treated, and BEV-treated endothelial monolayers. ICAM-1 fluorescence intensity per cell was quantified in HUVECs under static (0 dyn/cm^2^), low-flow (LFR; 1.5 dyn/cm^2^), and high-flow (HFR; 10 dyn/cm^2^) conditions. **(A)** Untreated HUVECs. **(B)** LPS-treated HUVECs (10 ng/mL). **(C)** HUVECs treated with *E. coli*-derived BEVs (1 × 10⁸ particles/mL). In A–C, each point represents the mean of three imaging fields from one independent microfluidic chip (n = 3 chips per condition); error bars represent mean ± SD. Comparisons in A–C were made using ordinary one-way ANOVA with Tukey’s multiple-comparisons test on the independent chip means. **(D)** Representative micrographs showing ICAM-1 immunofluorescence (green) and DAPI-stained nuclei (blue). **(E)** Violin plots of all nine experimental conditions displaying individual imaging-field measurements (nine fields from three chips per condition) to illustrate spatial and within-chip variation. Colors identify chips (chip 1, black; chip 2, blue; chip 3, red), and symbols identify imaging regions (triangle, inlet; circle, middle; square, outlet). Comparisons among all nine experimental conditions reported in the Results were performed using chip means (n = 3 per condition) with ordinary one-way ANOVA followed by Tukey’s multiple-comparisons test; the field-level points in E are displayed descriptively rather than treated as independent biological replicates. *p < 0.05, **p < 0.01, ****p < 0.0001.

Mean ICAM-1 fluorescence per cell was 0.097 arbitrary units (a.u.) under static conditions and 0.082 a.u. at LFR. This modest decrease under LFR was not statistically significant in the chip-level analysis (adjusted p = 0.1378). In contrast, the HFR mean was 0.455 a.u., approximately 4.7-fold higher than static and 5.6-fold higher than LFR (adjusted p < 0.0001 for both comparisons; **Fig. 3A**). These findings demonstrate that basal endothelial ICAM-1 expression varied with shear magnitude, with the greatest fluorescence observed under HFR conditions.

### LPS-associated ICAM-1 expression was greatest under high shear stress

LPS (10 ng/mL) was included as a bacterial inflammatory comparator to evaluate whether shear-dependent changes in endothelial ICAM-1 expression also occurred following exposure to a non-vesicular bacterial stimulus **(Fig. 3B,D)**.

Among LPS-treated HUVECs, mean ICAM-1 fluorescence per cell was 0.137 a.u. under static conditions, 0.175 a.u. at LFR, and 0.550 a.u. at HFR. The HFR mean was approximately 4.0-fold and 3.1-fold higher than the static and LFR means, respectively. Chip-level comparisons showed that HFR was significantly higher than both static and LFR (adjusted p < 0.0001 for both comparisons), while static and LFR did not differ significantly (adjusted p = 0.2433; **Fig. 3B**).

To distinguish treatment-associated responses from the effects of shear stress alone, LPS-treated cells were compared with shear-matched untreated controls using the independent chip means **(Fig. 3E)**. LPS treatment increased mean ICAM-1 fluorescence under static (0.137 versus 0.097 a.u.), LFR (0.175 versus 0.082 a.u.), and HFR (0.550 versus 0.455 a.u.) conditions. In the nine-condition chip-level comparison, LPS treatment significantly increased ICAM-1 fluorescence relative to shear-matched untreated controls under LFR (adjusted p = 0.0021) and HFR (adjusted p = 0.0016), but not under static conditions.

### BEV-treated endothelial cells exhibited elevated ICAM-1 expression under flow

To evaluate the influence of shear stress on BEV-associated endothelial activation, HUVECs were exposed to *E. coli*-derived BEVs (1 × 10^8^ particles/mL) under static, LFR, and HFR conditions **(Fig. 3C,D)**. Mean ICAM-1 fluorescence per cell increased from 0.164 a.u. under static conditions to 0.556 a.u. at LFR and 0.691 a.u. at HFR. Compared with static BEV-treated cells, fluorescence was approximately 3.4-fold higher at LFR and 4.2-fold higher at HFR (adjusted p < 0.0001 for both comparisons). Chip-level analysis also identified a significant increase from LFR to HFR (adjusted p = 0.0026; **Fig. 3C**).

BEV-treated cells had higher mean ICAM-1 fluorescence than their corresponding untreated controls in static culture (adjusted *p* = 0.0383) and at both shear levels (adjusted *p* < 0.0001 for each comparison). At 1.5 dyn/cm^2^, mean fluorescence per cell was 0.082 a.u. in untreated cells and 0.556 a.u. in BEV-treated cells, a 6.8-fold difference. At 10 dyn/cm^2^, the corresponding means were 0.455 and 0.691 a.u., a 1.5-fold difference. In chip-level comparisons of all nine conditions, BEV-treated cells exhibited significantly higher mean ICAM-1 fluorescence than LPS-treated cells under both LFR and HFR (adjusted p < 0.0001 for both comparisons), but not under static conditions.

Together, these findings demonstrate that BEV-associated ICAM-1 fluorescence depends on the mechanical environment: the greatest absolute mean occurred under high shear, whereas the largest relative difference from the shear-matched untreated mean occurred under low shear. The individual fields shown in **Fig. 3E** did not reveal an obvious, consistent inlet-to-outlet pattern on visual inspection, although position was not tested formally and the data did not demonstrate uniform shear stress.

### Cell-associated CMO fluorescence was higher under high shear stress than in static culture

We measured cell-associated CellMask Orange (CMO) fluorescence after exposing HUVECs to CMO-labeled BEVs under static, low-shear, and high-shear conditions (**Fig. 4A**). Mean fluorescence per cell was 0.0162 a.u. in static culture, 0.0781 a.u. at low shear, and 0.1472 a.u. at high shear (**Fig. 4C**). Fluorescence differed across conditions (exact Kruskal–Wallis test, *p* = 0.0036). In pairwise comparisons, fluorescence was higher under high shear than in static culture (adjusted *p* = 0.0219). Neither comparison involving low shear reached significance.

**Figure 4.**
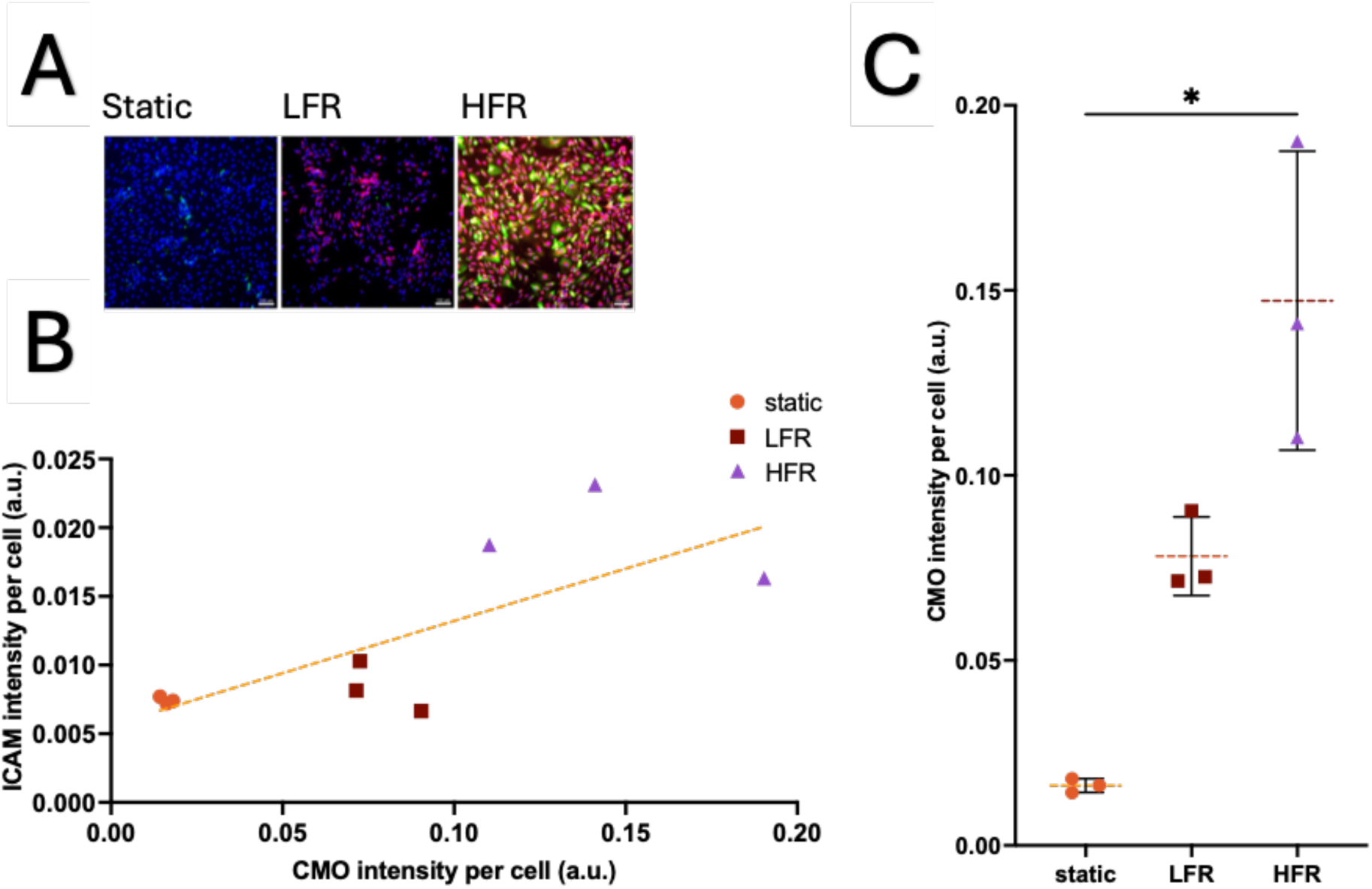
Shear-dependent changes in cell-associated fluorescence from CMO-labeled bacterial extracellular vesicles. HUVEC monolayers were exposed to CellMask Orange (CMO)-labeled E. coli-derived BEVs (1 × 10^8^ particles/mL) under static (0 dyn/cm^2^), low-shear (LFR; 1.5 dyn/cm^2^), and high-shear (HFR; 10 dyn/cm^2^) conditions. **(A)** Representative fluorescence micrographs showing DAPI-stained nuclei (blue), CMO-associated fluorescence (magenta), and ICAM-1 immunofluorescence (green). Scale bars are shown in the individual images. **(B)** Scatter plot comparing CMO fluorescence intensity per cell with ICAM-1 fluorescence intensity per cell using paired chip-level mean measurements (n = 9 across three shear conditions). Symbols distinguish static (circles), LFR (squares), and HFR (triangles). The dashed line represents the linear regression across all nine chip-level observations (R^2^ = 0.571, p = 0.0186). **(C)** Quantification of cell-associated CMO fluorescence intensity per cell across shear conditions. Each point represents the mean of three imaging fields from one independent microfluidic chip (n = 3 chips per condition). Horizontal lines and error bars represent mean ± standard deviation (SD). An exact Kruskal–Wallis test identified an overall difference among shear conditions (p = 0.0036). Dunn’s multiple-comparisons test identified a significant difference between static and HFR (adjusted p = 0.0219), but not between static and LFR or between LFR and HFR (adjusted p = 0.5391 for both). *p < 0.05. Fluorescence intensity is expressed in arbitrary units (a.u.).

Across the nine chips, mean cell-associated CMO fluorescence was positively associated with mean ICAM-1 fluorescence (R^2^ = 0.571, *p* = 0.0186, **Fig. 4B**). The regression pooled chips from all three shear conditions and therefore does not establish whether the association holds within a given condition.

## DISCUSSION

We have previously studied the effects of *E. coli*-derived BEVs on endothelial cells in static culture. Since endothelial cells are exposed to hemodynamic forces that shape their phenotype and their response to circulating inflammatory stimuli [22], we wanted to determine whether HUVECs would respond differently to BEVs under flow. We tested how defined laminar shear stresses change endothelial ICAM-1 expression after exposure to *E. coli*-derived BEVs. High shear raised baseline ICAM-1 in untreated cells. BEVs increased ICAM-1 relative to shear-matched controls under all three conditions, and the largest relative increase occurred at low shear, even though the highest absolute ICAM-1 occurred at high shear. BEVs also produced greater ICAM-1 than LPS under both flow conditions but not under static conditions. In exploratory experiments with fluorescently labeled BEVs, cell-associated fluorescence was higher under high shear than under static conditions. These results indicate that the mechanical environment affects how endothelial cells respond to BEVs, so studies done only in static culture may miss part of that response.

The clearest basal effect was higher ICAM-1 in untreated cells at 10 dyn/cm^2^ than under static or low-shear conditions. Mean ICAM-1 at 1.5 dyn/cm^2^ was slightly lower than static, but this difference was not significant in the chip-level analysis. We did not test the pathways that might explain it, such as Krüppel-like factor 2 (KLF2) and KLF4, which have been implicated in responses to laminar flow [23]. The high-shear result agrees with earlier reports that shear stress regulates endothelial adhesion molecules, including ICAM-1, in a time- and magnitude-dependent way [23]. Nagel et al. found that laminar shear selectively increased ICAM-1 in human endothelial cells without an inflammatory agonist [24], and other studies saw ICAM-1 rise along a spatial gradient of shear while VCAM-1 and E-selectin responded differently [22], [25]. Because laminar flow is usually linked to endothelial homeostasis, higher ICAM-1 under high shear should not automatically be read as dysfunction. KLF2, KLF4, nuclear factor-κB (NF-κB), and cytoskeletal mechanotransduction regulate different aspects of endothelial function and can give different outputs depending on shear magnitude, exposure time, and cellular context [26]. We interpret our result more modestly: shear sets different baseline levels of an adhesion molecule that takes part in vascular inflammation.

Shear magnitude also changed the response to BEVs. ICAM-1 in BEV-treated cells rose from static to low to high shear, with the highest absolute value at 10 dyn/cm^2^. Relative to untreated controls, however, BEVs raised ICAM-1 about 6.8-fold at low shear and 1.5-fold at high shear, so the high baseline at 10 dyn/cm^2^ accounts for part of why the added BEV effect looks smaller there. This extends our earlier static work, in which *E. coli*-derived BEVs induced dose-dependent ICAM-1 expression that was attenuated by Toll-like receptor 4 (TLR4) inhibition [3]. One explanation is that shear conditioning puts endothelial cells into different signaling states. Mechanotransduction involves the glycocalyx, cell junctions, integrins, the cytoskeleton, and transcriptional regulators, several of which also take part in inflammatory signaling [27]. We did not measure TLR4 activation, NF-κB nuclear translocation, or other mechanosensitive intermediates, so the mechanism remains open.

BEVs and LPS also behaved differently. BEVs increased ICAM-1 significantly over untreated controls under all three conditions, while LPS did so only under flow, and BEV-treated cells had higher ICAM-1 than LPS-treated cells under both flow conditions but not under static conditions. This fits Kim et al. [20], who found that E. coli-derived outer membrane vesicles increased endothelial adhesion molecule expression, including ICAM-1, more strongly than equivalent amounts of purified LPS. They also examined leukocyte adhesion under flow, whereas we compared how different levels of laminar shear affect ICAM-1 after BEV and LPS exposure.

Several differences between the stimuli could explain the pattern. BEVs carry membrane-associated LPS along with proteins, lipoproteins, and nucleic acids, and free LPS can reach the TLR4-MD-2 receptor complex through LPS-binding protein and CD14 [29], whereas intact vesicles may engage the endothelial surface differently. BEV and LPS doses were defined by different measures, so they should not be read as equivalent amounts of bacterial material, and our data cannot say whether the differences reflect endothelial responsiveness, delivery of the stimulus, or other shear-dependent processes.

Because ICAM-1 alone cannot separate a change in vesicle association from a change in endothelial responsiveness, we added exploratory experiments with CellMask Orange (CMO)-labeled BEVs. These lacked the controls of the ICAM-1 experiments, so we treat them as hypothesis-generating. Mean cell-associated CMO fluorescence was higher under both flow conditions than under static culture and highest at 10 dyn/cm^2^, but only the static versus high-shear difference was significant in the chip-level analysis. Fluorescence and ICAM-1 were positively related across chips, but this pooled relationship may partly reflect differences between shear groups and does not show that more BEV association causes more ICAM-1. Our highest fluorescence at high shear differs from studies of mammalian extracellular vesicles, which reported enhanced uptake under low or disturbed shear [16], [28].

These results matter for how BEVs are studied in sepsis. Endothelial dysfunction is a major part of sepsis pathophysiology, contributing to leukocyte recruitment, microvascular permeability, dysregulated coagulation, and impaired tissue perfusion [30], and endothelial cells in different parts of the circulation experience different flows. Identical bulk BEV concentrations produced different ICAM-1 responses under static, low-shear, and high-shear conditions, so static experiments alone may not capture how BEVs interact with the endothelium in flowing blood.

Our shear conditions were meant to represent distinct mechanical environments, not the full hemodynamics of one vascular bed, but they give a basis for testing how local flow shapes vascular responses to microbial particles and how mechanical and biochemical signals combine to activate the endothelium.

This study has several limitations. Epifluorescence cannot distinguish vesicles bound to the apical membrane or glycocalyx from internalized ones, and confocal orthogonal sectioning, transmission electron microscopy, or membrane-impermeable quenching assays would resolve BEV endocytosis versus surface retention under flow [31]. CMO fluorescence also may not reflect the number of intact vesicles, since membrane dye transfer and residual dye-associated material could contribute. We used an endothelial monoculture under laminar flow, whereas vessels have complex hemodynamics, spatial shear gradients, and constant crosstalk with immune and perivascular cells; multicellular models and non-uniform or disturbed flow would show where vessels are most vulnerable during sepsis. Measuring glycocalyx components such as heparan sulfate and hyaluronic acid under different shear conditions, and modifying the glycocalyx enzymatically, would show whether it acts as a barrier to BEVs or helps retain them near the endothelial surface. Statistically, we used three independent chips per condition, with three imaging fields per chip averaged within each chip and the chip means used as independent observations. The nine-field distributions in Fig. 3E show within-chip variability, but we did not formally test position-specific differences, so their absence cannot establish fully developed or uniform flow. Finally, the chip-level correlation between CMO and ICAM-1 fluorescence may reflect differences among shear groups rather than an association within each condition.

## CONCLUSION

Shear stress changed both baseline endothelial ICAM-1 and the response to E. coli-derived BEVs. High shear raised baseline ICAM-1, BEVs increased ICAM-1 over shear-matched controls under every condition, and LPS did so only under flow. The highest absolute ICAM-1 occurred under high shear, while the largest relative BEV effect occurred under low shear.

Exploratory CMO experiments revealed greater cell-associated fluorescence under high shear, suggesting shear-dependent differences in BEV-associated material at the endothelial surface. Vascular MPS and other in vitro models of endothelial–BEV interactions should therefore include defined levels of shear stress. Future work that separates vesicle transport, surface association, and internalization from endothelial mechanosensitivity should clarify how these responses arise during bacterial infection.

## ACKNOWLEDGEMENTS

Research reported in this manuscript was supported by the National Institutes of General Medical Sciences (NIGMS) of the National Institutes of Health under grant number R35GM153461 to Thomas Gaborski. The content is solely the responsibility of the authors and does not necessarily represent the official views of the National Institutes of Health.

## Notes

### Competing Interest Statement

The authors have declared no competing interest.

